# Discovery of novel compounds as potent activators of Sirt3

**DOI:** 10.1101/2022.01.05.475007

**Authors:** Célina Reverdy, Gaetan Gitton, Xiangying Guan, Indranil Adhya, Rama Krishna Dumpati, Samir Roy, Santu Chall, Gauthier Errasti, Thomas Delacroix, Raj Chakrabarti

## Abstract

Among the sirtuin enzymes, Sirt3 is one of the most important deacetylases as it regulates acetylation levels in mitochondria, which are linked to the metabolism of multiple organs and therefore involved in many types of age-related human diseases such as cancer, heart diseases and metabolic diseases. Given the dearth of direct activators of Sirt3, the identification of new modulators could be a key step in the development of new therapeutics. Here we report the discovery of Sirt3 modulators, including activators, through the use of DNA encoded library technology (DEL). The most enriched compounds after DEL selection against SIRT3 were evaluated according to their activity and affinity. Our best activator seems at least as potent as Honokiol (HKL) while the docking studies suggest that our modulators interact with Sirt3 at an atypical site. Our results establish the attractiveness of the DEL technology in identifying novel and potent Sirt3 activators and, therefore, in associated therapeutic applications.

## 1 Introduction

Sirtuins are a family of proteins known as Class III histone deacetylases. These NAD^+^ dependent enzymes are highly conserved in mammals throughout evolution.^1^ They deacetylate various substrates and control pivotal processes in the cells from genomic integration to stress response and metabolism.^2^ There are 7 sirtuins in mammals that differ in their localization and function. Sirt1, 6 and 7 are in the nucleus and are involved in DNA maintenance and transcription, while Sirt2, located in the cytoplasm, contributes to tubulin polymerization.^3,4^ Sirt3, 4 and 5 are localized in the mitochondria out of which Sirt3 is the prime deacetylase in regulating metabolism and promoting adaptation to nutrient starvation.^5^ Sirt3 is a key regulator of mitochondrial metabolism including the tricarboxylic acid cycle (TCA), urea cycle, amino acid metabolism, electron transport chain or oxidative phosphorylation, ROS detoxification and mitochondrial unfolded protein response. Sirt3 is predominantly expressed in tissues rich in mitochondria such as kidney, heart, brain and liver tissue. Thus, Sirt3 activity in these tissues is crucial to maintain mitochondrial function. Sirt3 has been shown to regulate aging, neurodegeneration, liver, kidney, heart diseases and other metabolic diseases.^6^ Due to its diverse role in the biological functioning of the cells, it has emerged to be a promising therapeutic target and several studies over the years have reported some activators or inhibitors of Sirt3.^6^ The catalytic core of Sirt3 is highly conserved and consists of a large Rossmann-fold domain and a smaller zinc binding domain.^7^

Sirt3 is a NAD dependent lysine deacylase, requiring the cofactor NAD^+^ to remove acyl groups from lysine groups of its substrates.^8^ The acetylated substrate and the NAD^+^ cofactor bind in between the two domains forming a stabilized cofactor binding loop. In the active conformation, the NAD^+^ is buried in the C-pocket facilitating the nicotinamide (NAM) release and an alkylamide intermediate formation. Further hydrolysis of the intermediate yields the deacetylated peptide product and 2’-*O*-acetyl-ADP-ribose (OAADPr).^9^

Most sirtuin modulators that have been described to date lack specificity and potency.^10^ The most widely known Sirt3 activator, Honokiol, has shown some therapeutic effects in heart disease, cancer and metabolic diseases but the mechanism remains to be elucidated. Recently, Upadhyay *et al*.^11^ reports a non-allosteric mechanism of activation of Sirt3 by Honokiol. 3-TYP, a known Sirt3 inhibitor, is used only as a probe showing no therapeutic application.^6^

Thus, developing effective Sirt3 modulators, and in particular activators, remains an attractive field of research. There is a high demand to identify specific and high affinity Sirt3 modulators showing a significant therapeutic activity. Herein, we report the discovery of Sirt3 modulators, including potent activators, using an affinity-based selection thanks to DNA encoded library technology (DEL). Our results establish the attractiveness of the DEL technology in identifying novel and potent Sirt3 activators and, therefore, in associated therapeutic applications.

## 2 Results and discussion

### 2.1. DNA encoded library selection

In our effort to discover novel small molecules that modulate sirtuin activities, we used DNA encoded library technology (DEL),^12^ a robust hit identification approach that uses large collections of diverse DNA-encoded small molecule libraries which are screened for their affinity against a desired protein target. The technology provides an efficient method to select a broad chemical space of structures. It is also an attractive strategy as it requires negligible amounts of target protein to do the selection experiments and it identifies ligands regardless of their functional activity. In the past few years, DEL has been successfully used to identify hits against several soluble targets.^3^ It has also been used against sirtuins^14–16^, but not for the discovery of activators.

We produced and screened a 3880-member DNA encoded library based on the design defined by Mannocci *et al*.^17^ and consisting of the coupling of 20 amino acids with 194 carboxylic acids. This library was chosen because of the relatively high molecular surface area of the compounds, which can span larger binding sites potentially overlapping with the enzyme active site, as well as contribute enhanced binding affinity. The affinity mediated selection against Sirt3 was performed both under the presence of Carba-NAD (a stable NAD analog) and the co-product OAADPR to identify compounds that differentially bind to Sirt3 under such conditions. The Carba-NAD or OAADPr along with the acetylated or deacetylated peptide substrate is necessary to produce functional Sirt3 conformations.^18,19^

For each screening, a no target screening was concurrently performed that served as the negative control. This enabled us to discard all the compounds in the library that bound non-specifically to the nickel beads. All the screening experiments were performed at 4 °C to retain Sirt3 activity. During the selection procedure, special care was taken to wash the beads post incubation of the protein and the library to get rid of non-specific binders as much as possible. Only one round of selection was performed, and final eluates were amplified, clustered and sequenced by Illumina iseq100. Prior to selection, we sequenced the native library to ensure that all the compounds are present at nearly similar concentrations so that the selection is not biased against any compound which might be present at a higher concentration than others. Fig. 1 shows the 3D plot of the native library showing the reads of the DNA barcodes corresponding to each compound. This is representative of the distribution of the compounds in the library. We observed that all the 194 derivatives of each of the 20 amino acids showed a similar distribution in terms of their frequency post next generation sequencing. Only AA5 derivatives appeared to be slightly overrepresented in the native library. The initial frequencies of the compounds helped us to normalize the final output and cancel out any bias that might occur due to difference in the concentrations of the individual compounds in the native library.

**Fig. 1.**
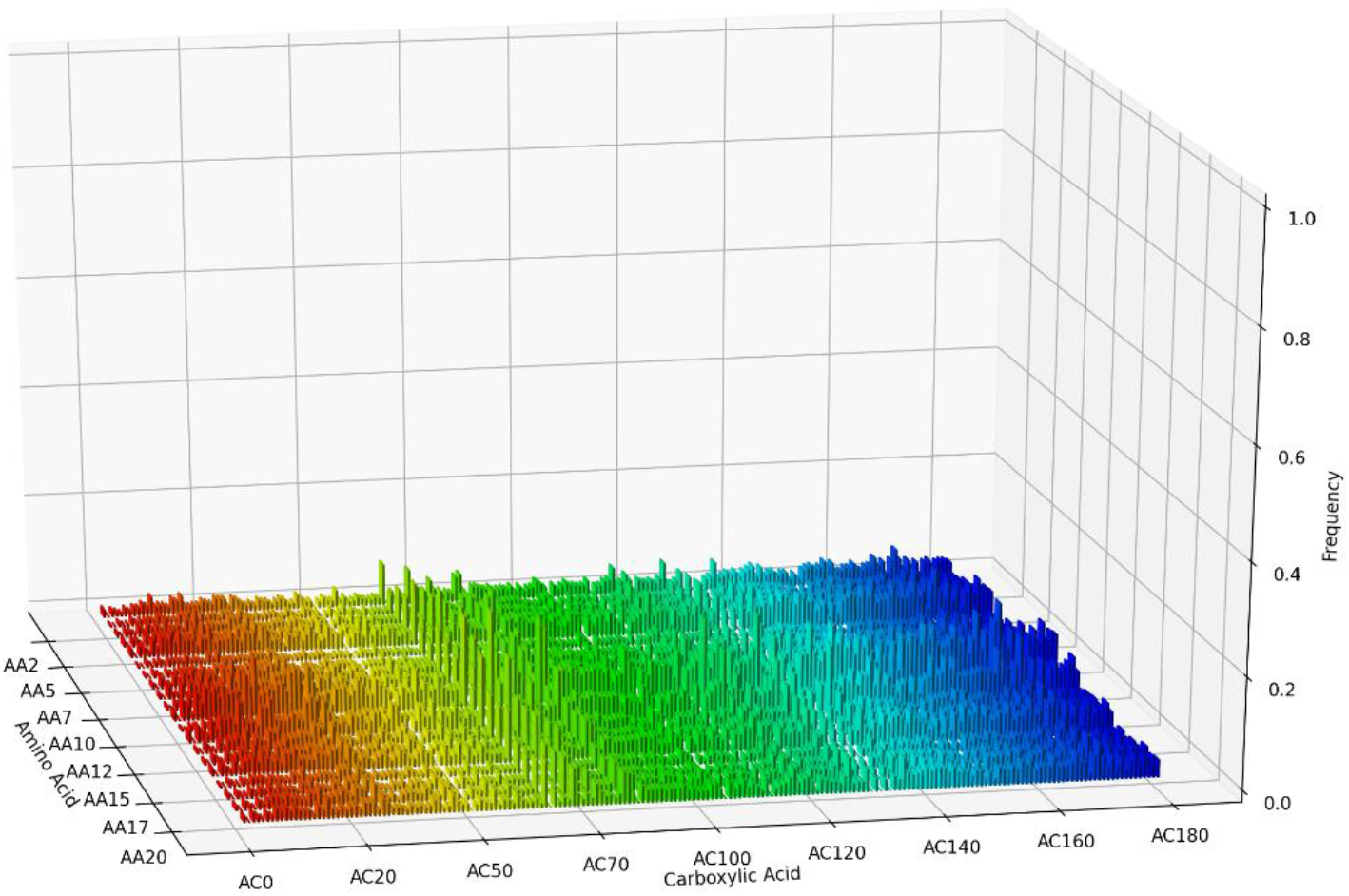
Frequency of all the compounds in the native library. All the derivatives of the 20 amino acids show a similar distribution over its 194 carboxylic acid counterparts in the library. AA5 appeared to be slightly overrepresented

#### 3880-member DNA encoded library selection against Sirt3

A first selection experiment was done without any target (negative selection): we observed that none of the compounds in the library showed any affinity to the nickel beads confirming the wise choice of the immobilization matrix.

Fig. 2 and 3 show the results of the selection against the Sirt3 with co-factors (Carba-NAD and acetylated peptide in Fig.2; OAADPr and deacetylated peptide in Fig. 3). The enrichments of the compounds over the background (no target selection) are pictured after normalizing their abundance with respect to the total number of DNA sequence reads obtained for each selection. The top 10 highly enriched samples were selected for characterization, represented in Table 1. All these compounds showed an enrichment between 12-16 fold over the no target control selection.

**Fig. 2.**
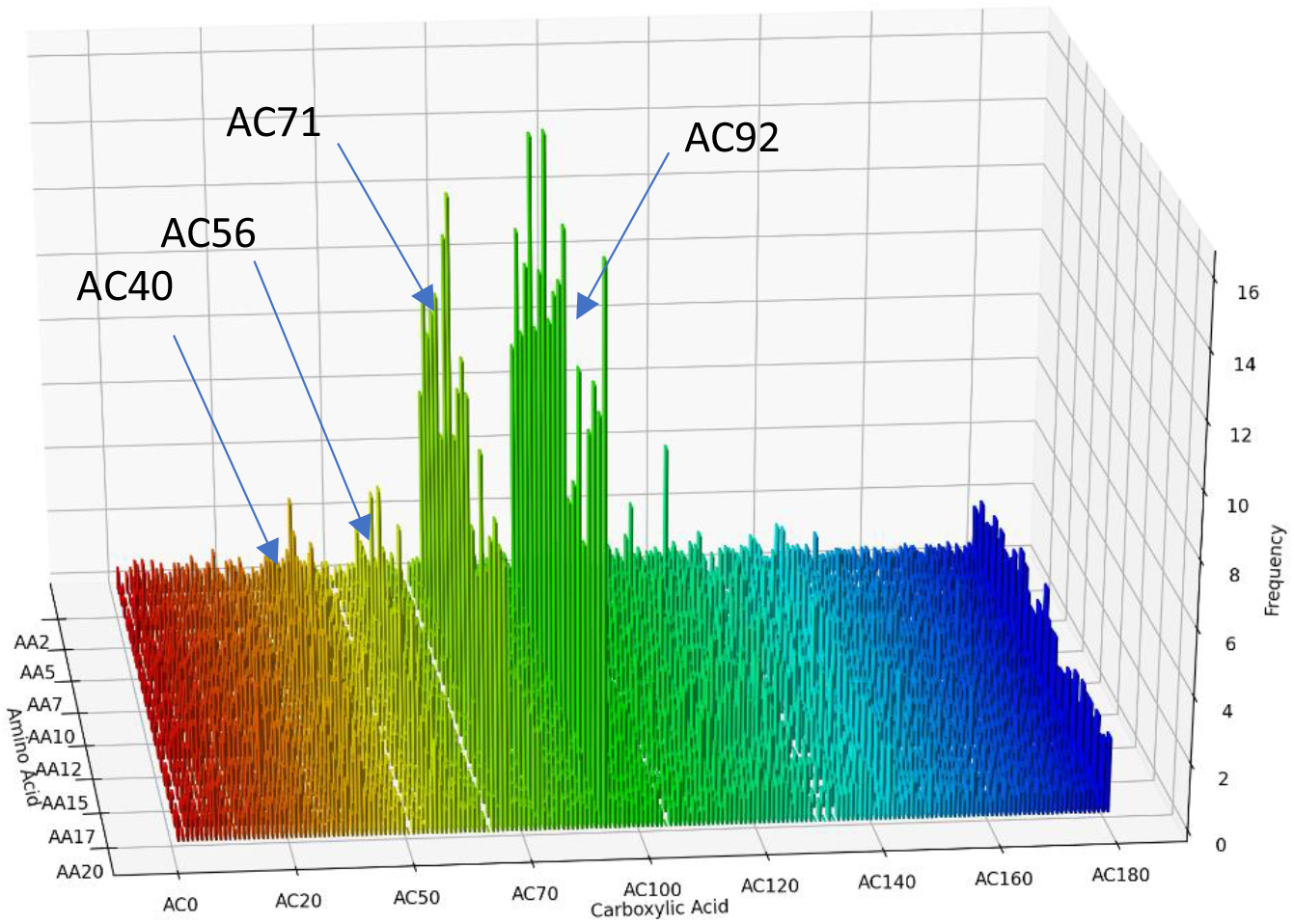
Sirt3 selection in the presence of Carba-NAD and acetylated peptide. The derivatives of AC71 and AC92 show the highest enrichment. Among the other, few derivatives of AC40 and 56 also show significant enrichment. The Y-axis represents the enrichment calculated for each compound as the ratio of their frequencies in the target selection to their frequencies in the no target selection.

**Fig. 3.**
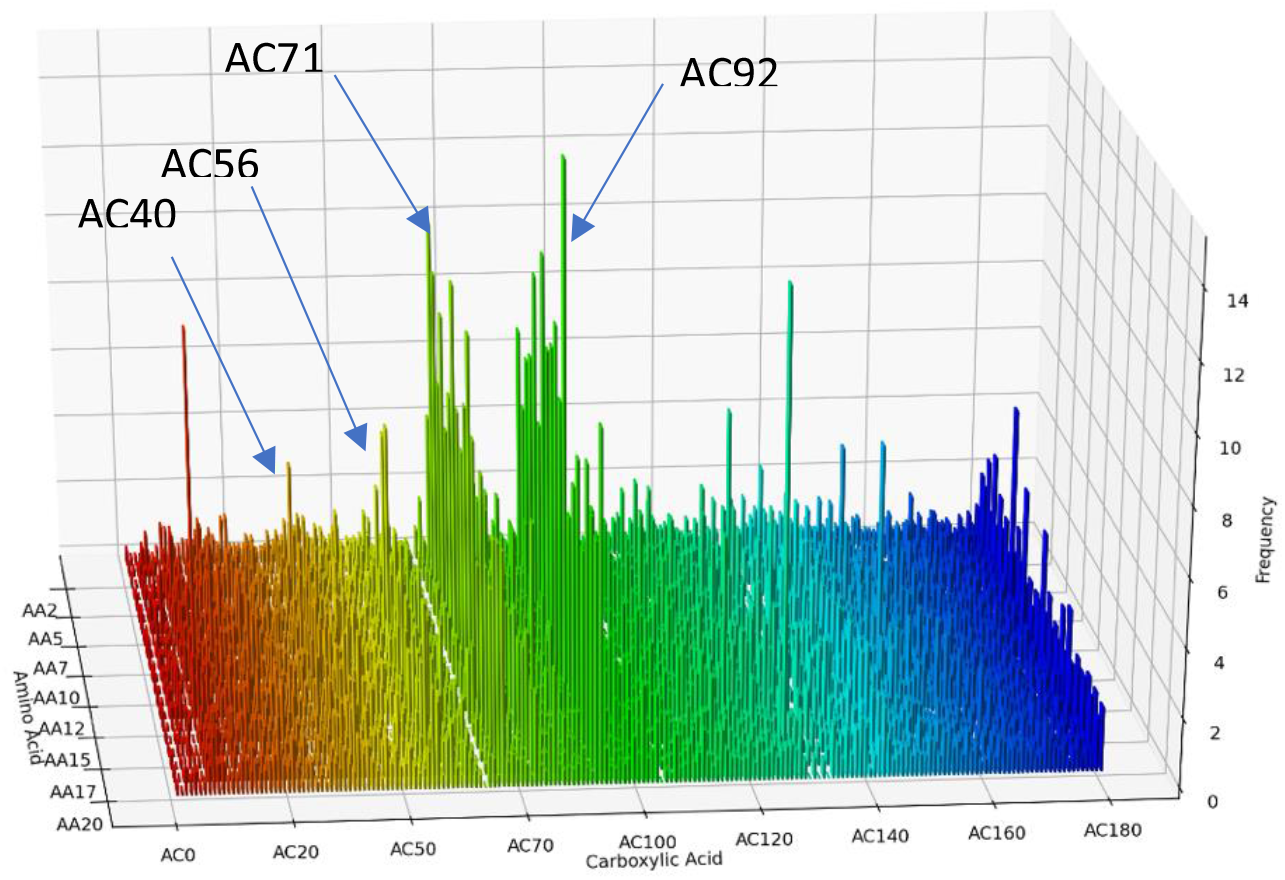
Sirt3 selection in the presence of OAADPr and deacetylated peptide. The derivatives of AC71 and AC92 show the highest enrichment. Among the other, few derivatives of AC40 and AC56 also show significant enrichment. The Y-axis represents the enrichment calculated for each compound as the ratio of their frequencies in the target selection to their frequencies in the no target selection.

**Table 1.**
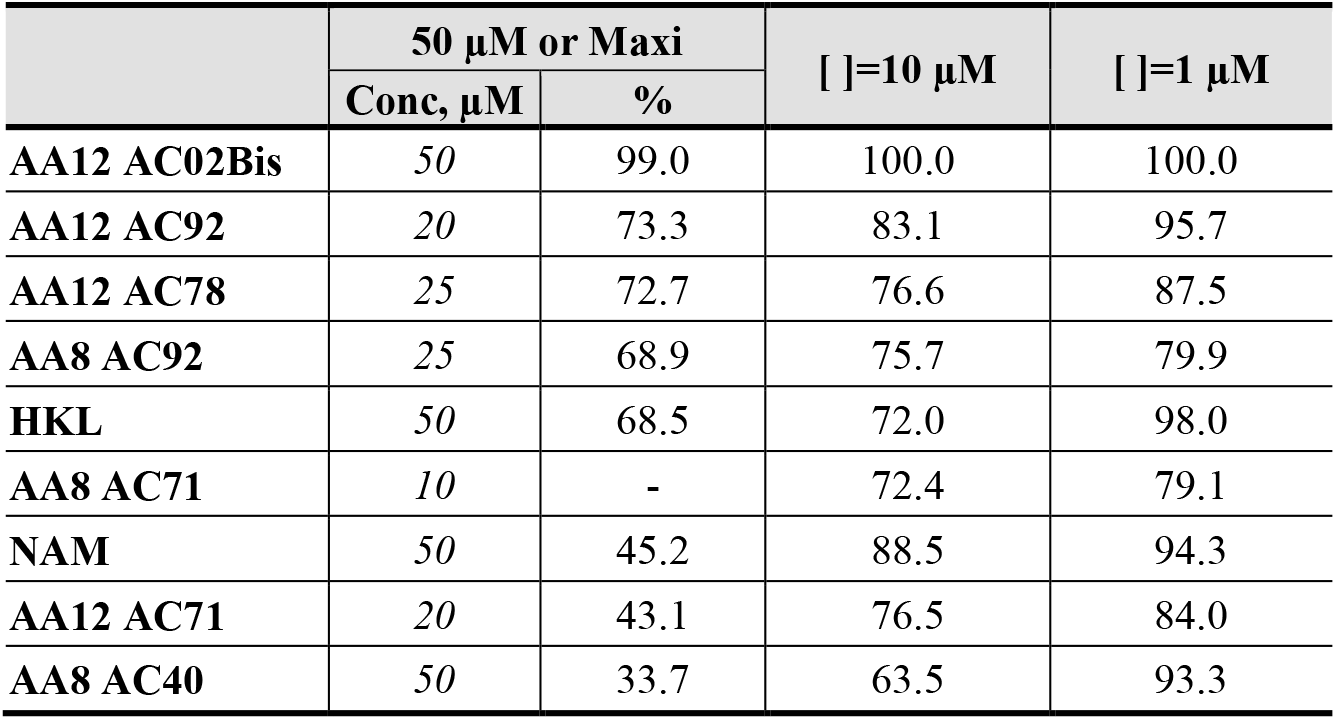
The potency of the selected compounds on Sirt3 deacetylation activity under steady state conditions.

Interestingly, for both the selection conditions, in the presence of Carba-NAD and OAADPr, the same derivatives of carboxylic acids, AC71 and AC92, showed the highest enrichment. Among others, a few derivatives of AC40 and AC56 showed significant enrichment between 4 and 8 fold over negative selection and would be worth investigating. The compounds were not overrepresented in the native library, further corroborating the hypothesis that enrichment was observed only due to binding to Sirt3. These compounds showed similar enrichment when performed in duplicates for both the selection conditions. This emphasizes the role of both the AC71 and AC92 derivatives as putative binding partners of Sirt3, which is our rationale behind choosing them for further analysis. All the top 10 hits are derivatives of either AC71 or AC92. Amino acids AA5, AA8, AA11 and AA12 are also common in the derivatives of both (Fig.4).

**Fig. 4.**
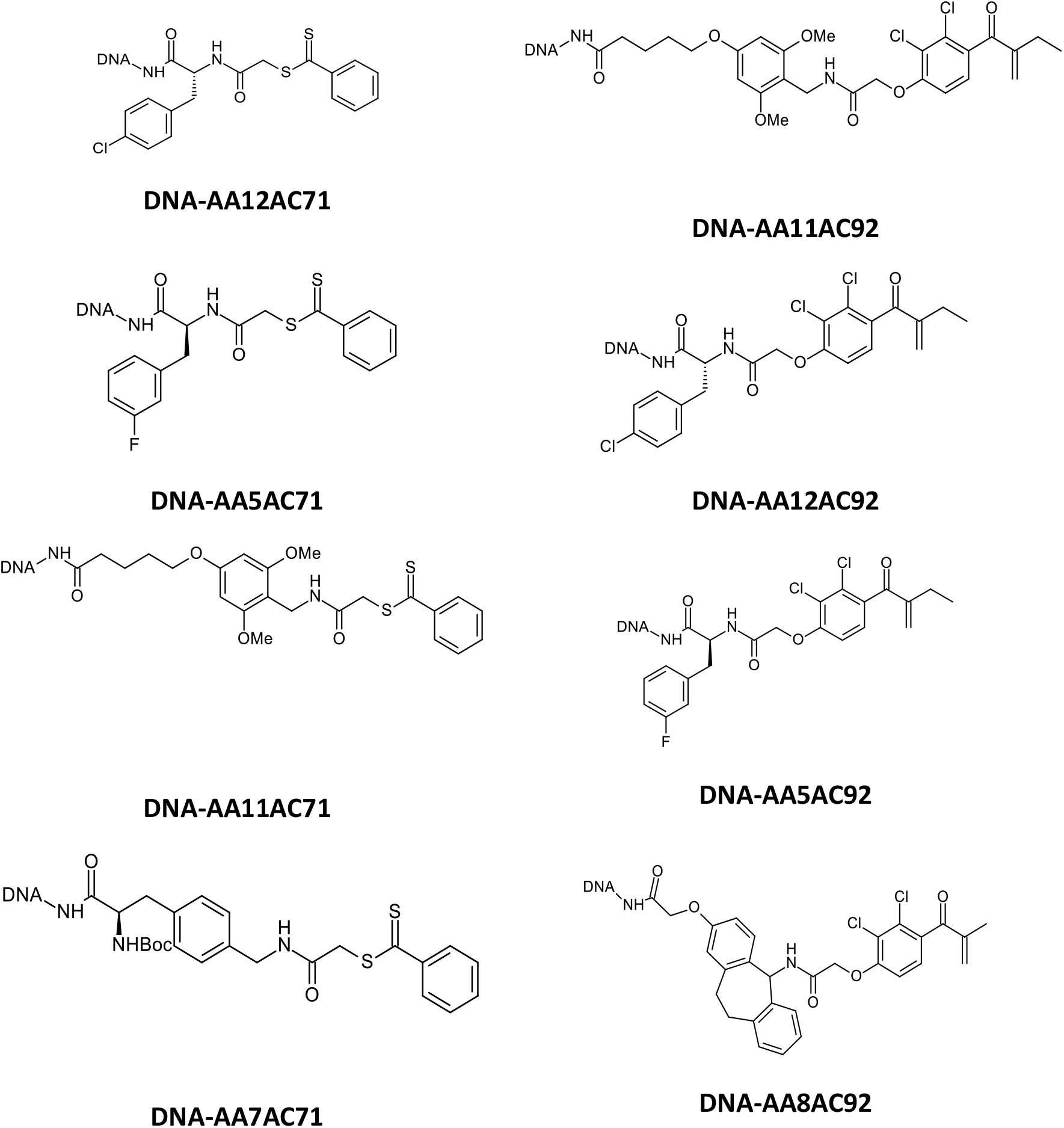
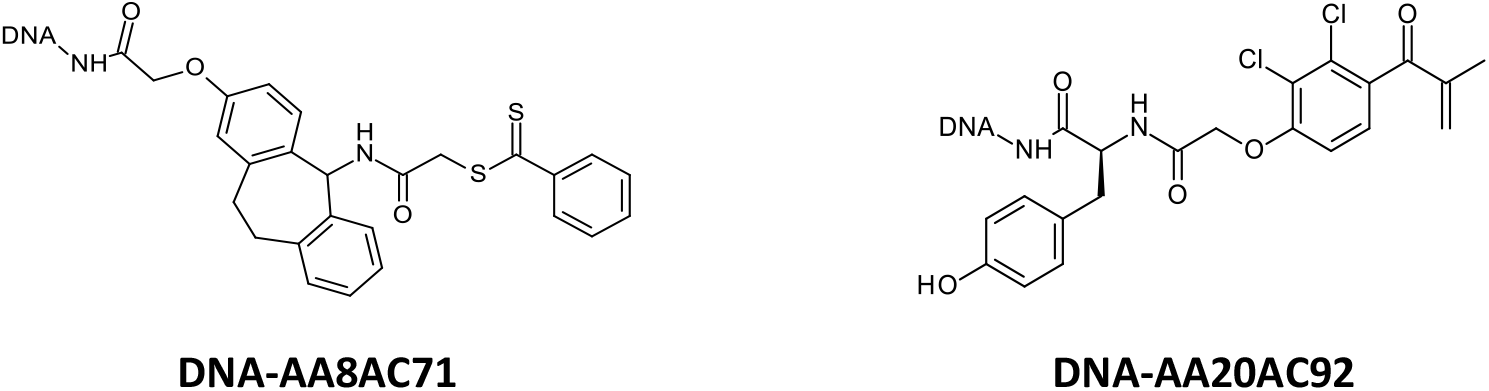
The chemical structures of the top 10 hits from both the Sirt3 selection experiments in the presence of Carba-NAD or OAADPr.

Thus, it was decided to synthesize 7 compounds among which 6 (AA12AC92, AA12AC78, AA12AC71, AA8AC92, AA8AC71, AA8AC40) were derivatives from AA8 and AA12 that showed the best enrichments, and one compound (AA12AC02Bis) that showed no enrichment, serving as a negative control (Fig. 5).

**Fig. 5.**
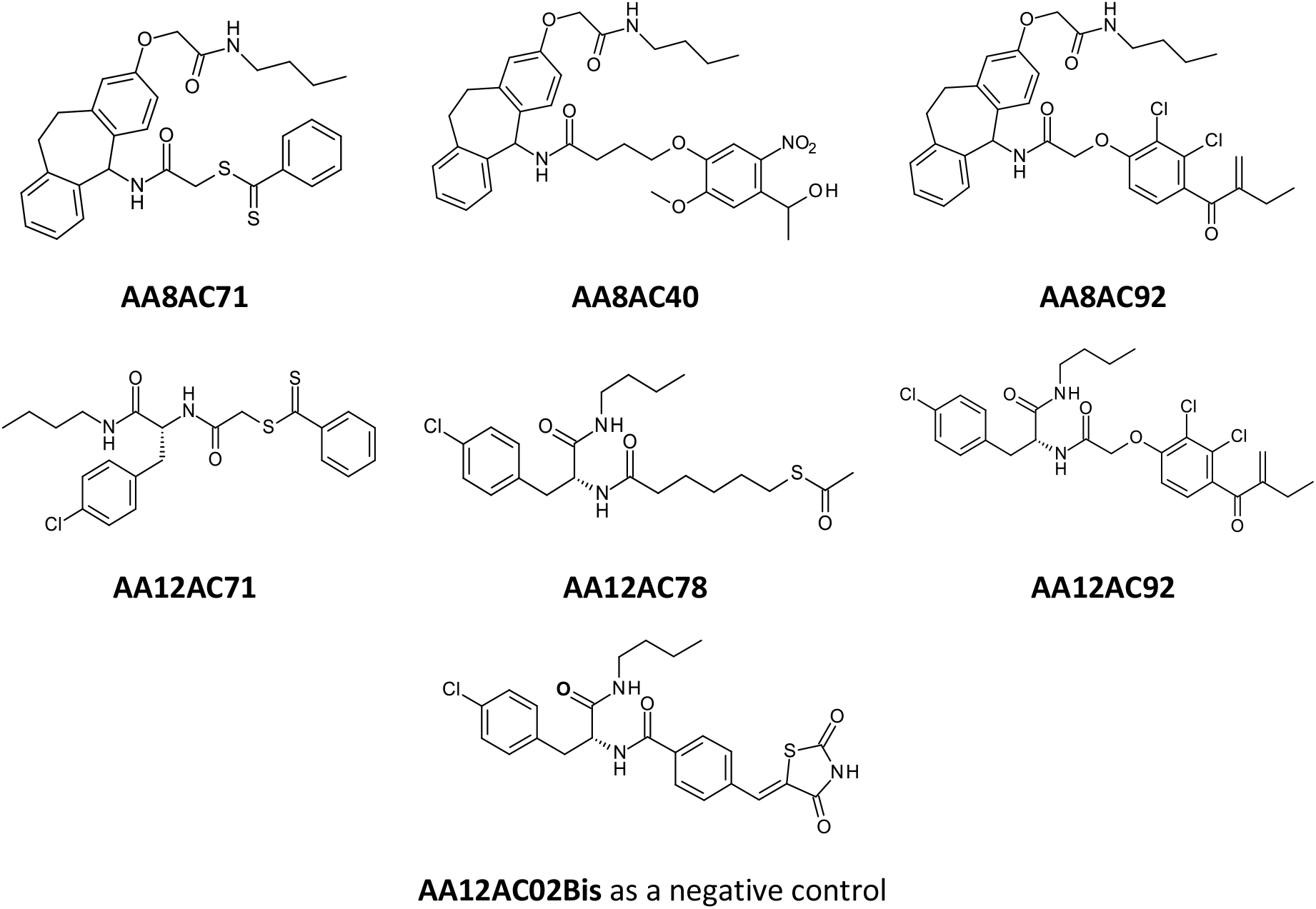
The chemical structures of the off-DNA top 6 hits (derivatives from AA8 and AA12) and one off-DNA negative control (AA12AC02Bis).

These compounds were synthesized off-DNA (The DNA chain has been replaced by a butyl group in order to test their modulation effect on Sirt3 by activity tests and to validate their binding affinity by switchSENSE® analyses.)

### 2.2. Activity assays

#### Sirt3 modulation effect of Hit Compounds

The 6 top ranked compounds from DEL screening were tested for their modulation effect of Sirt3 using fluorescence-based and HPLC-based assays. The effect of the compounds on enzymatic turnover is expected to experience three phases (pre-steady state, steady state, and post-steady state) during the enzymatic reaction^11^. The pre- and post-steady states can be considered as non-steady state, in which the enzyme concentration vs. OAADPR co-product concentration ratio is high and low, respectively. Depending on the initial ratio of enzyme to limiting substrate in the system, the net effect of the tested compounds over the course of the reaction can be either inhibiting or activating, with activation occurring if this ratio is above a threshold^11,20^.

#### Sirt3 modulation effect under steady-state conditions

The selected compounds show different degrees of inhibitory effect under the steady-state condition (Fig. 6). It was found that the Sirt3 deacetylation activity was inhibited by 66.3% in the presence of 50 µM compound AA8AC40. It was also found that the inhibition level increases as the compound dose increases. The IC_50_ values for the compounds are expected to fall in the 10-20µM range (Table 1). The negative control (AA12AC02Bis) has no contribution of Sirt3 modulation. The activity results agree with the DEL screening output.

**Fig. 6.**
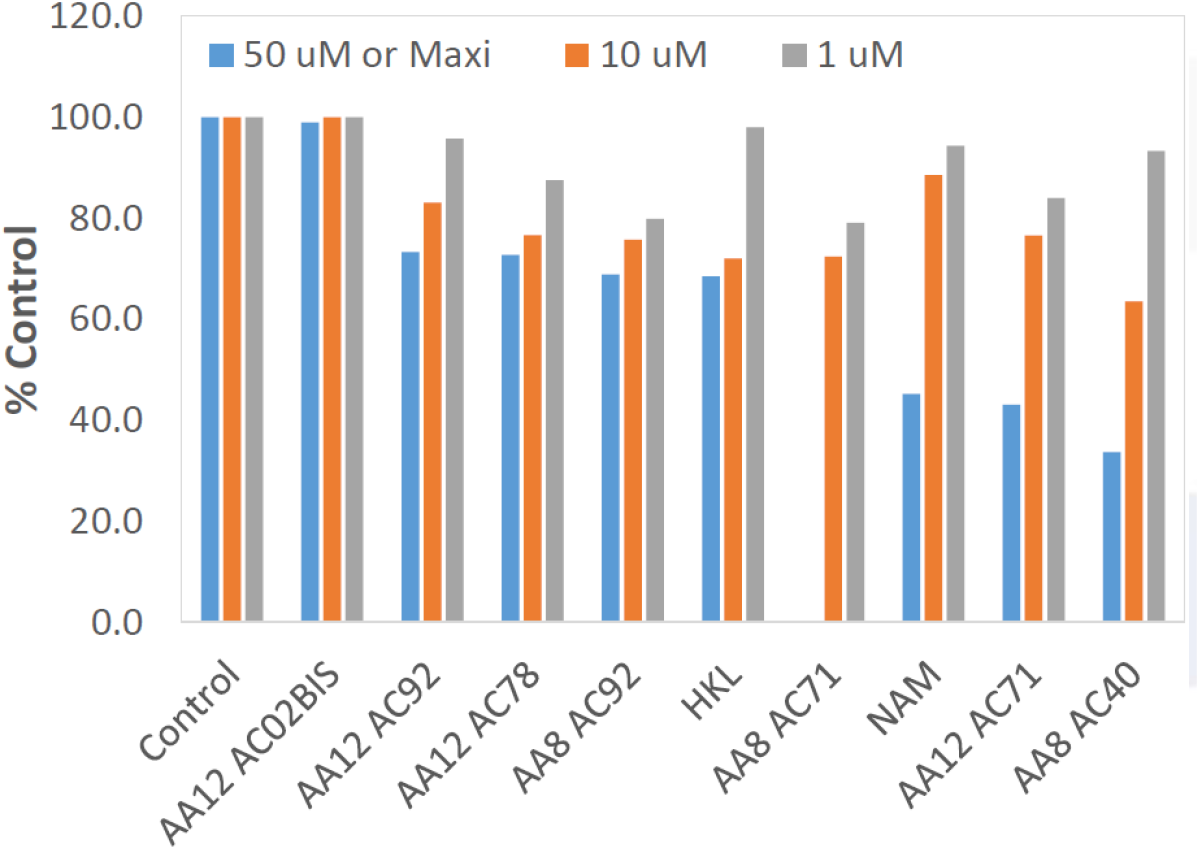
Effect of top ranked hit compounds on Sirt3 deacetylation activity under steady state conditions. Bar diagram of % control by different cpds ([NAD^+^] =1 mM, [FdL2 peptide] = 50 µM, [cpd] = 1, 10, 50 µM or maximum soluble concentration, [E]_0_/[NAD]_0_ = 0.000245, Time point=30 min, n=2).

#### Non-steady state activation of Sirt3

The activation of SIRT3 was detected under non-steady state condition in the presence of 3 out of 7 compounds, e.g. AA8AC40, AA12AC92, AA12AC71 (Fig.7). Compound AA8AC40 provides the most activating effect (Table 2), comparable to that of Honokiol (HKL), which was extensively studied as a non-steady state Sirt3 activator in our previous work.^20^ We note that for such non-steady state activators the extent of activation is predicted to be higher for a higher initial ratio of enzyme to limiting substrate concentration^20^; as such the activation values reported in Figs. 7 and Table 2 do not represent the maximum extent of activation possible with these compounds.

**Fig. 7.**
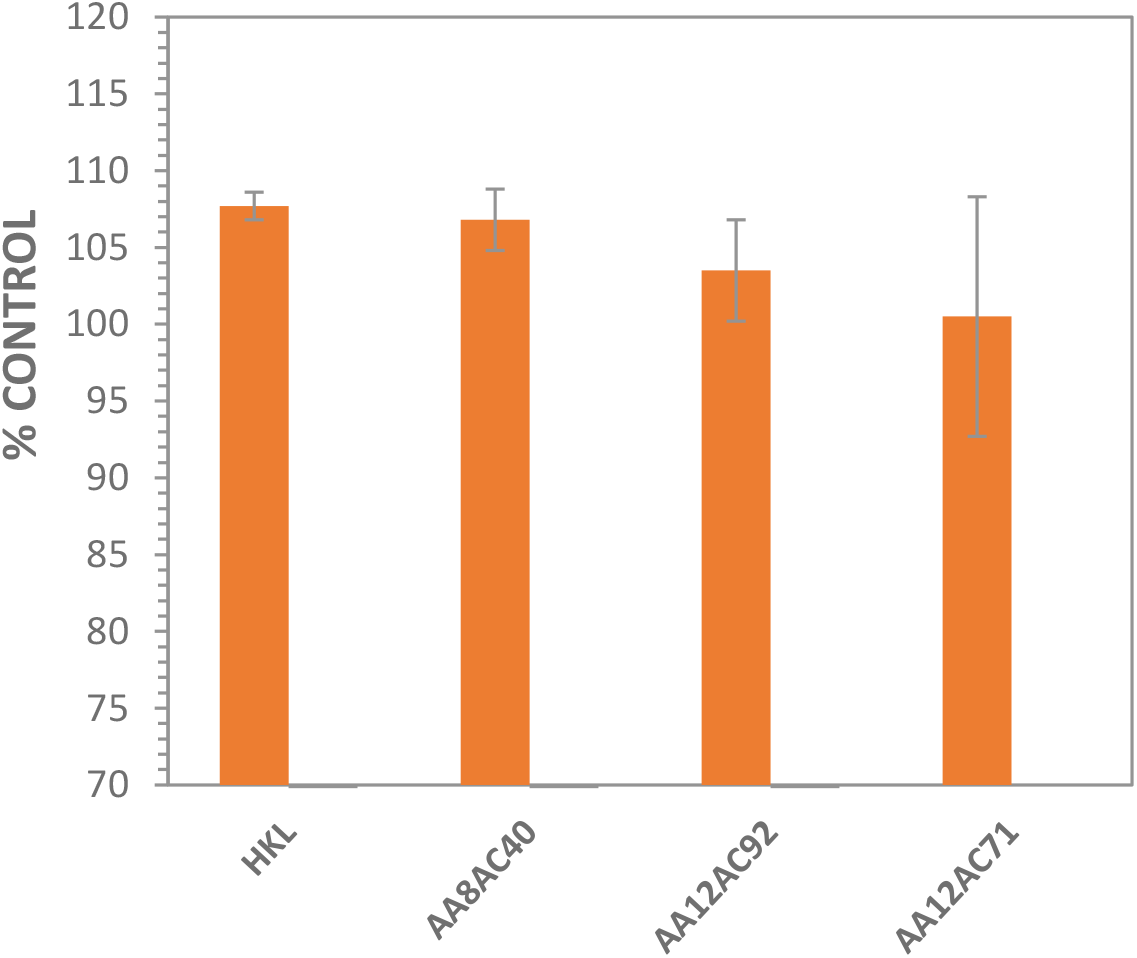
Sirt3 activation by top ranked hit compounds. Bar diagram of % control by different compounds ([NAD^+^] =50 µM, [MnSOD K122] = 600 µM, [cpd] = 10 µM or maxi, [E]_0_/[NAD]_0_ = 0.1223, Time point=2 min, n=3).

**Table 2.** The potency of the selected compounds on Sirt3 deacetylation activity under non-steady state condition.

| [modulator]=10 μM | % Control |
| --- | --- |
| **HKL** | 107.7 ±0.9 |
| **AA8AC40** | 106.8 ±2.0 |
| **AA12AC92** | 103.5 ±3.3 |
| **AA12AC71** | 100.5 ±7.8 |

### 2.3. Protein-compound docking

Ligand docking, as described in Materials and Methods, was performed on top-binding ligands from the DEL, as identified by the affinity mediated selection assay. However, the large size and flexibility of each DNA-tagged and combinatorially-generated ligand made it impossible to dock the entire complex to Sirt3. To address this problem, we decided to only dock the structure of combinatorial ligands, in their C-terminal amidated form, to Sirt3. In addition to most active combinatorial ligands, we also choose a few inactive ligands, also identified by the same assay, as negative controls to validate the choice of binding site and docking procedure. Structures for some positive and negative controls used in our docking studies are shown in Fig. 8.

**Fig. 8.**
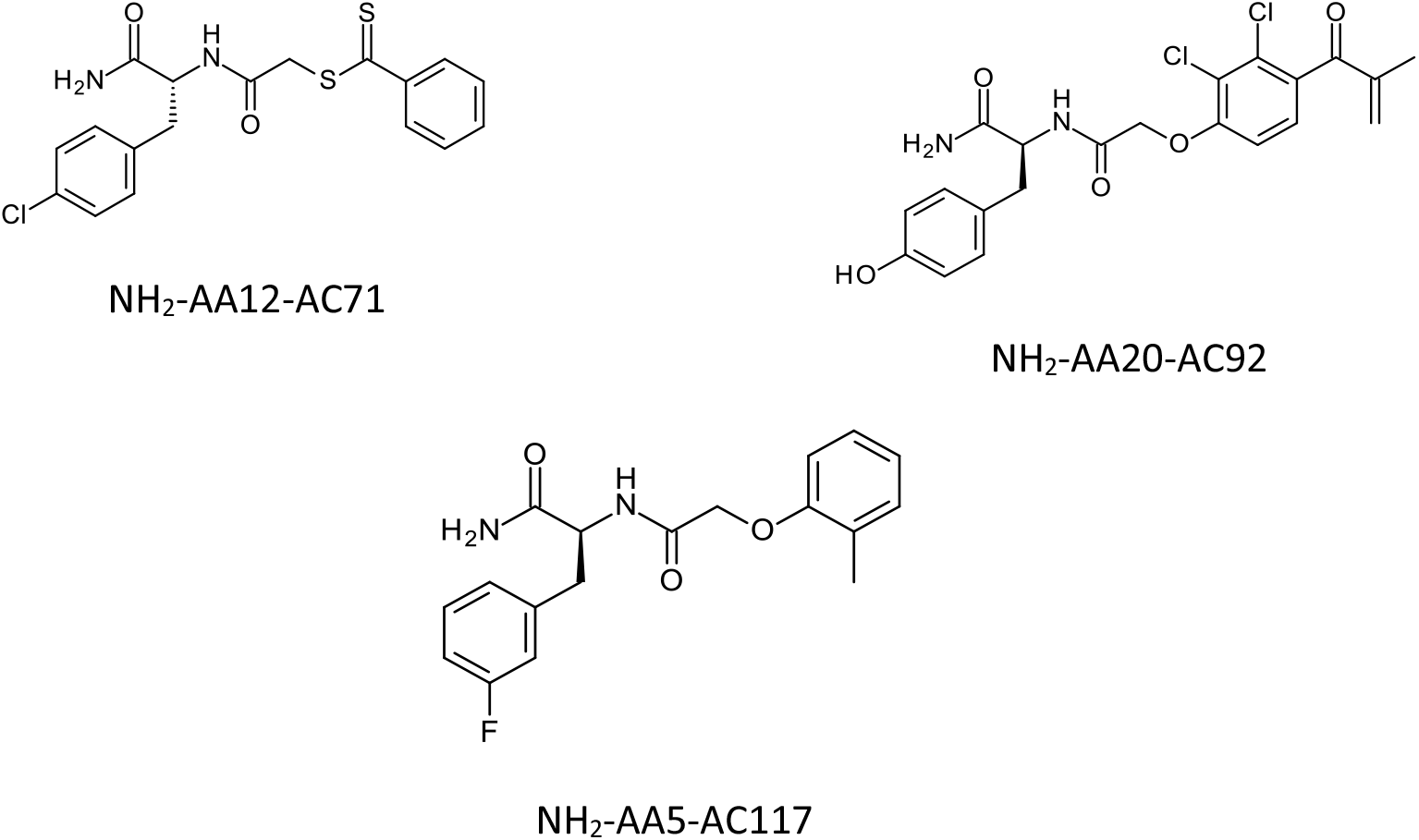
C-terminal amidated forms for selected active and inactive ligands identified by affinity selection assay. Structures of C-terminal amidated version of AA12AC71, an active ligand, AA20AC92, another active ligand, and AA5AC117, an inactive ligand. Compounds depicted were used for all ligand-docking studies.

Despite the substantial size differences between DEL-derived ligands and compounds known to bind at internal sites in Sirt3, such as Ex-527^19^ and Honokiol, we decided to first explore known internal sites in that protein, as shown in Fig. 9. However, our attempts at docking both the active and inactive hits from the DEL-library to internal sites in two structures of Sirt3, 4BVG and 4FVT, were not successful (not shown). These results were not unexpected for two reasons, which are as follows: (1) there is no worthwhile overlap between pharmacophores or other structural features of the hits obtained from the DEL-library and known compounds capable of binding at internal sites in peptide and NAD (or Carba-NAD) bound structures of Sirt3 such as Ex-527; (2) The hits, even without their DNA-tags, are noticeably longer and bigger than Ex-527 and the available space within internal sites, in the presence of peptide and Carba-NAD, is not adequate to properly accommodate ligands that are even slightly larger than Ex-527. Accommodating larger compounds in internal sites in those two structures would require the displacement of NAD, or Carba-NAD, as it is seen in complexes of Sirt3 with some potent inhibitors.^14^

**Fig. 9.**
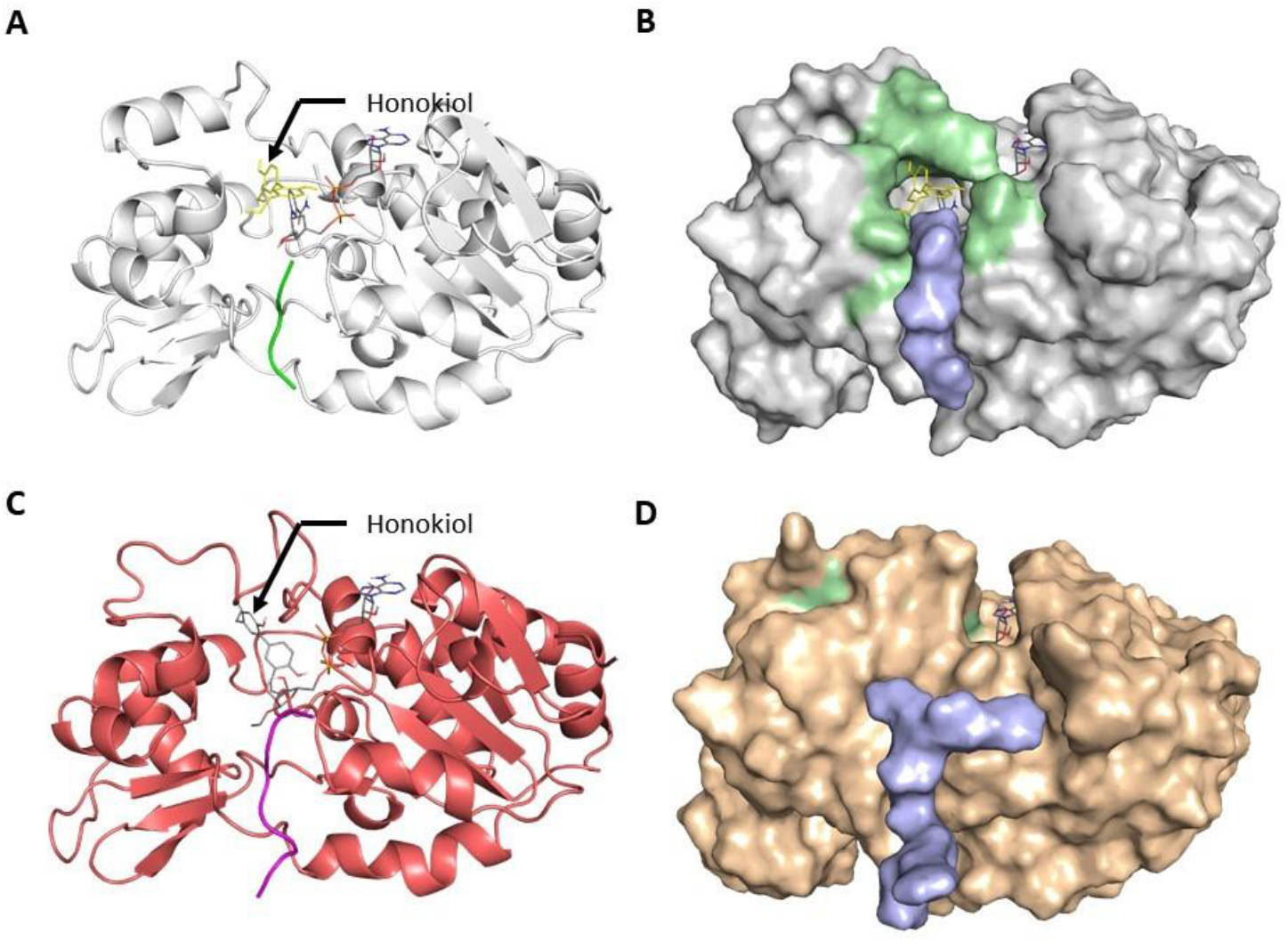
Location of internal binding site in 4FVT and 4BVG. (A) Cartoon rendering of 4FVT, (B) Molecular surface rendering of 4FVT with residues within 4.5 Å of docked Honokiol being highlighted with green, and ACS peptide as light blue. (C) Cartoon rendering of 4BVG and (D) Molecular surface rendering of 4BVG with residues within 4.5 Å of docked Honokiol highlighted with green and ACS peptide as light blue.

Attempted docking of hits and control at internal site within both SIRT structures (4FVT and 4BVG) revealed a few additional problems. First, clustering of top poses for hits is poor at internal sites within both structures, and a significant percentage of those poses lie partially outside the cavity (not shown). Second, the directionality of top docked poses for many hits is not compatible with location of DNA-tag linkage. Inactive controls from library also exhibit substantially better clustering than hits when docked to internal sites in both structures (not shown). Hence it is unlikely that hits from DEL-library bind at internal sites in Sirt3, leading us to the possibility that they bind to more exposed sites which are on (or near) the surface of protein.

We then started to search for putative binding sites in Sirt3 which were either atypical or external. 4FVT and 4BVG were scanned, using a geometry-based method in MOE, to locate potential binding sites. One atypical binding site large enough to partially accommodate positive library hits was identified in 4FVT. A similar scan of 4BVG revealed two more atypical potential binding sites large enough to accommodate these compounds. Attempts to dock positive hits to the atypical binding site in 4FVT revealed some degree of overlap between top docked poses for a few of the positive hits,; however, the degree of pharmacophore overlap for top poses of different ligands was still poor and no conserved pattern of interactions with receptor was seen (not shown). The location of the first atypical potential binding site in 4BVG is shown in Fig. 10.

**Fig. 10.**
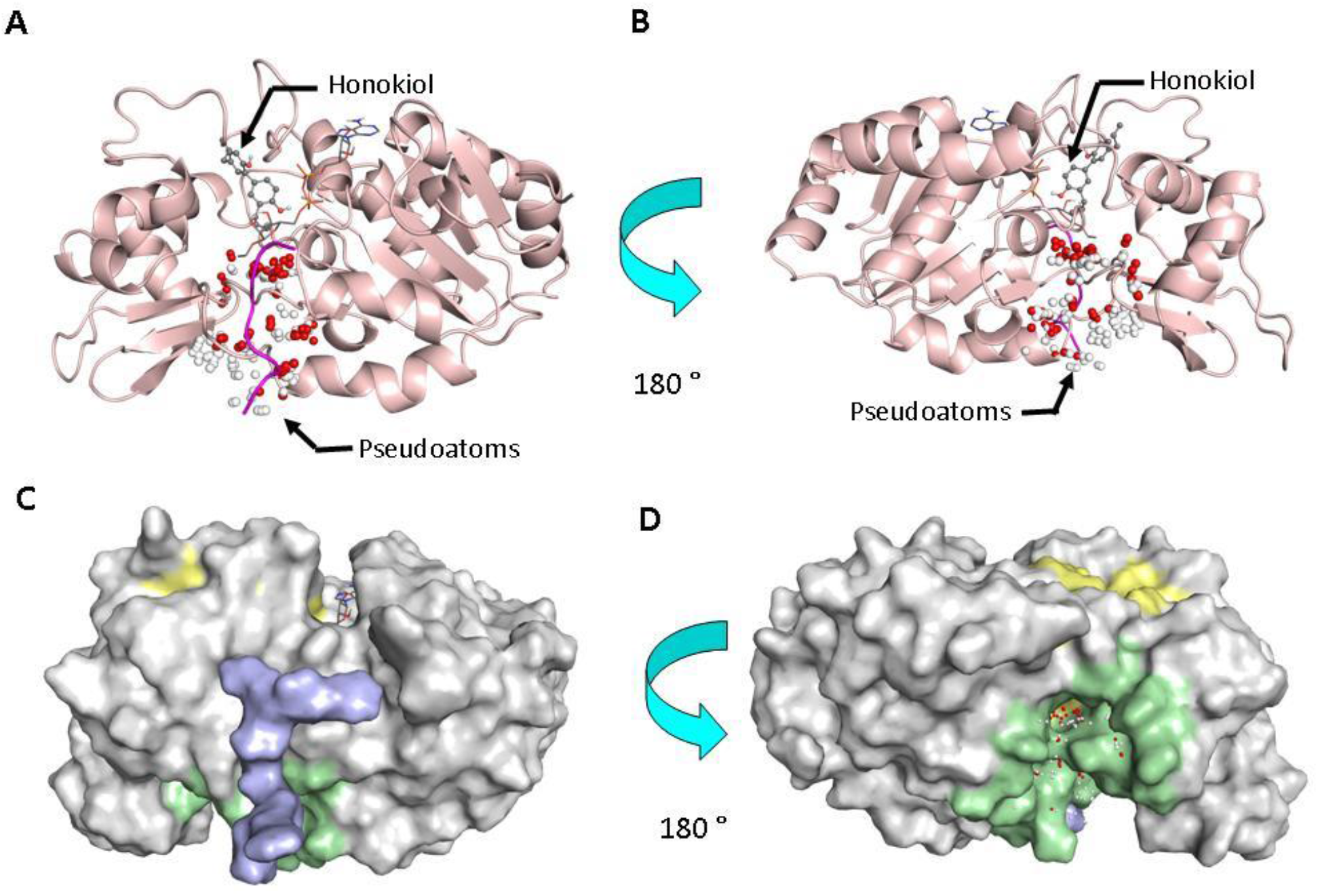
Location of the first and larger atypical site on surface of 4FVT. (A) Cartoon rendering showing relative position of Honokiol docked at internal site and pseudoatoms which highlight the position of atypical potential binding site in 4BVG, (B) Rotated version of model from A showing the relative location of docked Honokiol and pseudoatoms in 4BVG, (C) Surface rendering of 4BVG with residues within 4.5 Å of Honokiol highlighted with yellow and first atypical binding site using green, while ACS peptide is blue and (D) Rotated version of model from C showing another relative position of two sites from another viewpoint.

Docking at the first putative atypical binding site in 4BVG revealed a high degree of overlap for the top docked poses of most of the positive hits. Only 2/10 positive hits (AA8AC71, AA8AC92) did not consistently overlap with the top docked poses of the other 8/10 positive hits as shown in Fig. 11. Some other hits such as AA8AC40 did dock at the site, but with a different orientation.

**Fig. 11.**
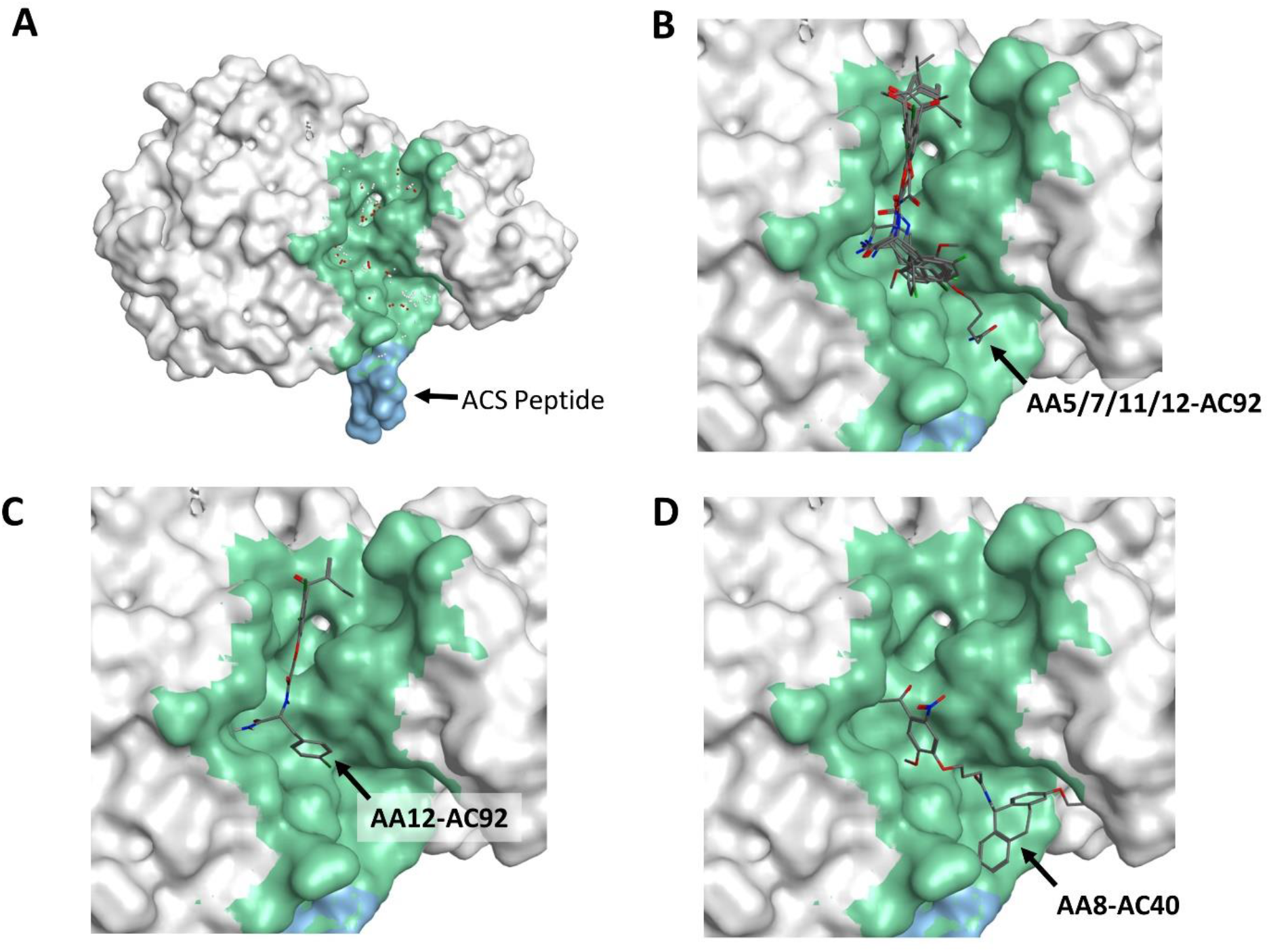
Location of Site 1 in 4BVG and top poses of docked hits. (A) Location of first putative atypical site, defined by pseudo-atoms, in 4BVG is almost antipodal to its well-known internal site. Residues lining this external site are highlighted with green, while ACS peptide using cyan. (B) Close-up of top docked poses of AA5/7/11/12-AC92 at the putative atypical site. (C) Close-up of top docked pose for AA12-AC92 at site. (D) Close-up of top pose of AA8-AC40 at putative site.

Top poses of most of the positive hits also demonstrated a high degree of pharmacophore overlap and conserved interactions with residues lining the site. Notably inactive controls also docked well at this site but displayed fewer interactions (hydrophobic and hydrophilic) than the hits with lining residues. The main hydrogen bond interactions of most active hits with site residues occur at A241, L244 and E246 and can be seen in Fig. 12. Even AA8AC40 interacts with E246 despite its different orientation (Fig. 13). Control compounds with low to zero activity dock to the receptor in a similar manner to active hits but are deficient in their ability to form the hydrogen bonds. Thus, even though hydrophobic interactions account for majority of receptor-ligand interactions, the two or three hydrogen bonds seems to determine whether a given hit is active or not.

**Fig. 12.**
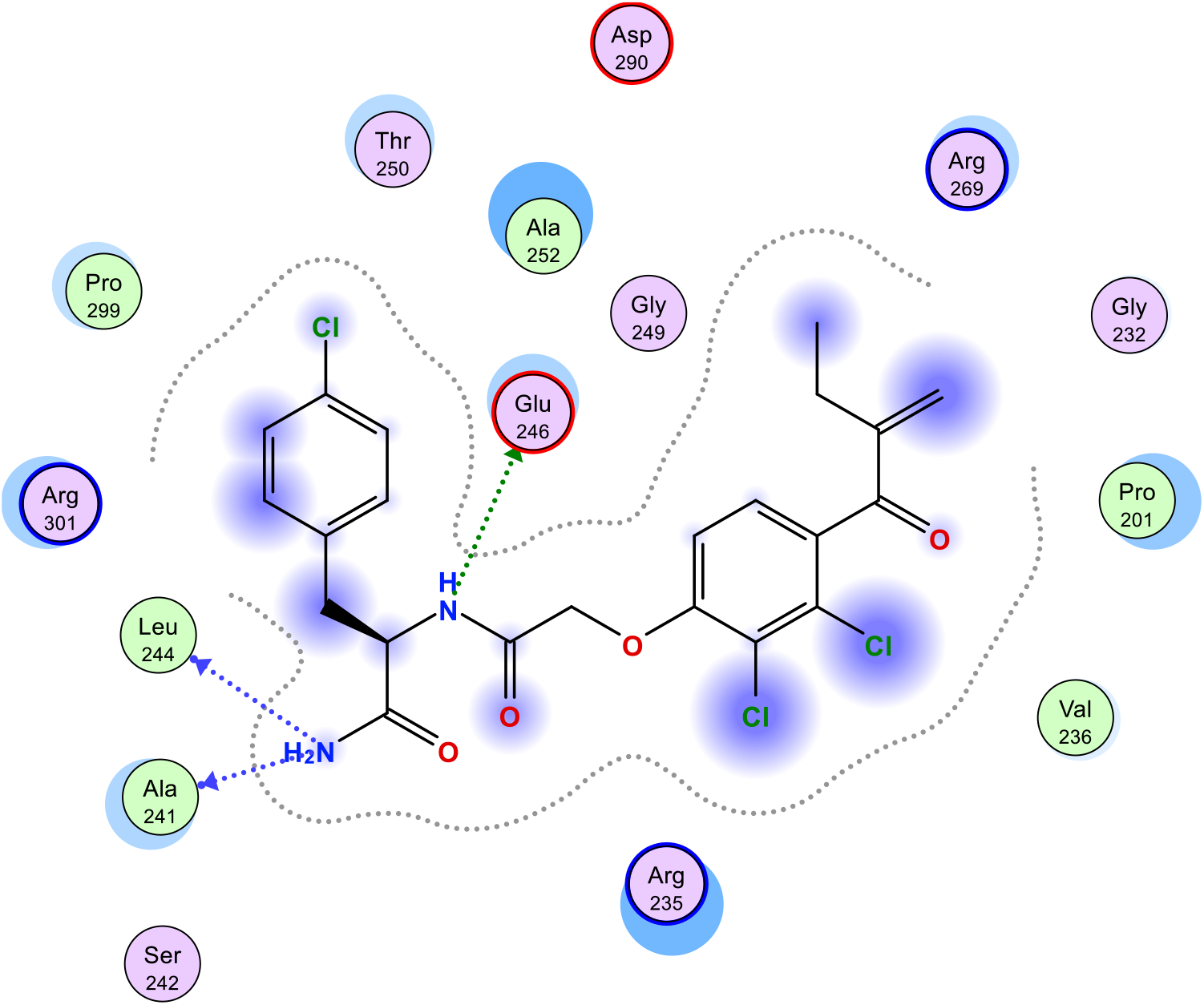
Hydrogen bonding and interaction pattern of one active hit (AA12-AC92) from DEL4000 library. The most important hydrophilic interactions seem to be between the compound and A242, L244 and E246. Note hydrophobic and cation-p interactions between aromatic rings in ligand and adjacent residues such as side chain of Leu244, Ala252, Val236, Arg301 and Arg235.

**Fig. 13.**
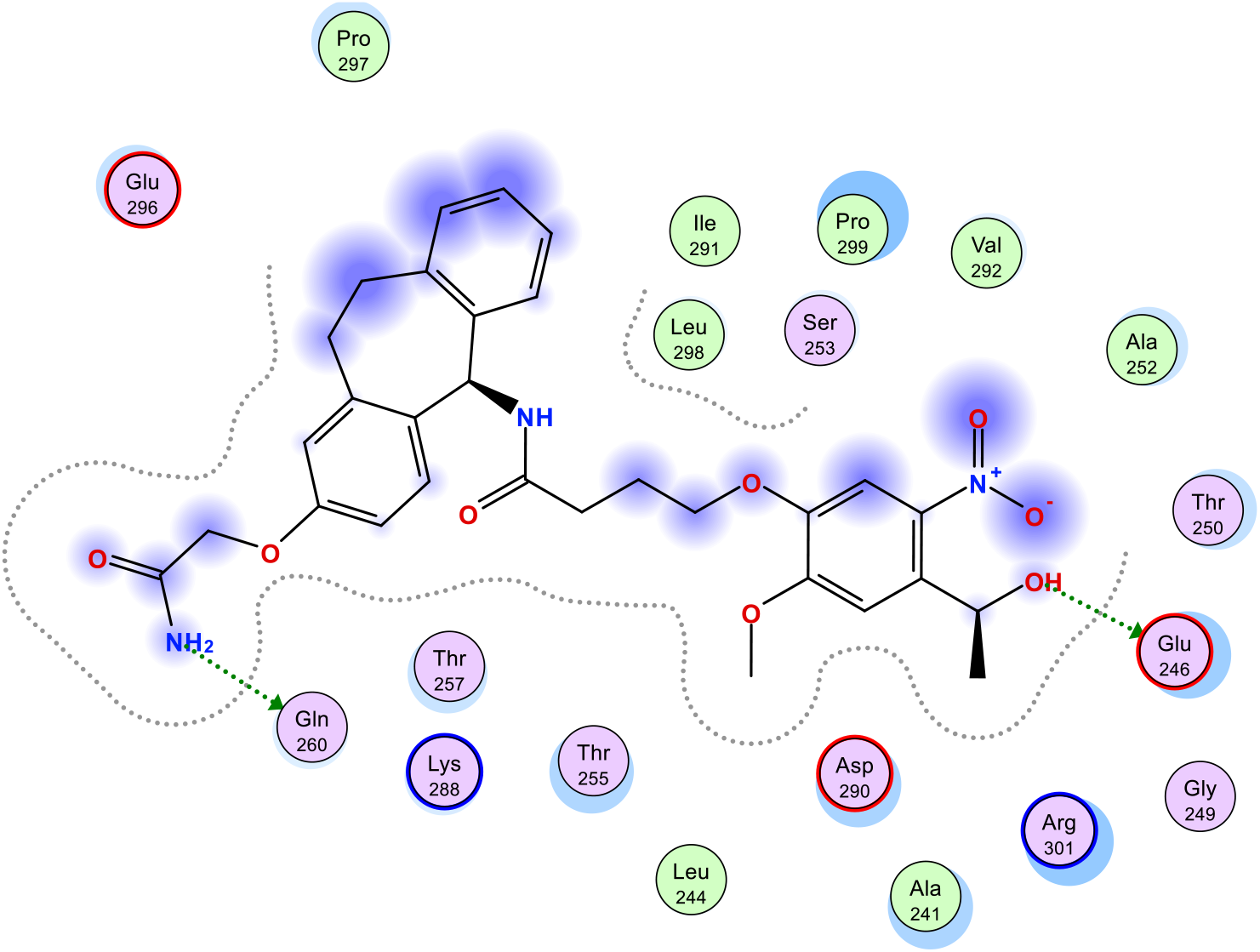
Hydrogen bonding and interaction pattern of one active hit (AA8-AC40) from DEL4000 library. The most important hydrophilic interactions seem to be between the compound and E246 and Q260. Note hydrophobic and cation-p interactions between aromatic rings in ligand and adjacent residues such as sidechain of Leu 298, Pro299 and Arg301.

A second putative atypical site (not shown) was identified, but the degree of pharmacophore overlap among the top poses of hits is nowhere close to that seen at the atypical Site 1 in 4BVG, and therefore the latter is the most likely binding site for the hits from the DEL-library. It should be noted that developing ligands for the atypical Site 1 in 4BVG might be harder than usual for low molecular weight small molecules, since it is significantly shallower and more exposed to the solvent than is typical for small-molecule sites. This further validates the choice of library design in the present work.

### 2.4. Binding Affinity Tests

Binding tests were performed on the top hits via SwitchSENSE®. It is based on short DNA nanolevers (48bp or 96bp) that can switch on 24 gold surface spots on a microfluidic chip. The switch of the DNA is mediated by alternating the voltage across the surface. The motion of the levers is tracked in real time (µs scale) through time-resolved single photon counting using a fluorescence probe signal present on one DNA stand. The complementary DNA strand can be cross-linked to a ligand through amine or thiol coupling or click-chemistry. Upon binding of an analyte, the hydrodynamic friction of the levers is first affected followed by the movement of the levers. This change is used by the system’s software to describe the absolute size/conformation of the ligand and complexes and to determine the kinetics of the interaction (k_on_, k_off_, K_d_).^21^

Affinity tests were performed with the compound AA8AC40 since it showed good Sirt3 activation effect under the non-steady state condition. Fig. 15 indicates that compound AA8AC40 binds to Sirt3 with a K_d_ value of 4.6 µ M (Table 3). Interestingly, in the presence of cofactors – OAADPr and deacetylated peptide-the binding affinity of AA8AC40 with co-structure does not change significantly, meaning the binding event still occurs. These results fall in agreement with the docking study presented above. Due to the size of the AA8AC40, it most likely binds partially within the C pocket of Sirt3 as well as the external surface of the Sirt3 enzyme.

**Fig. 15.**
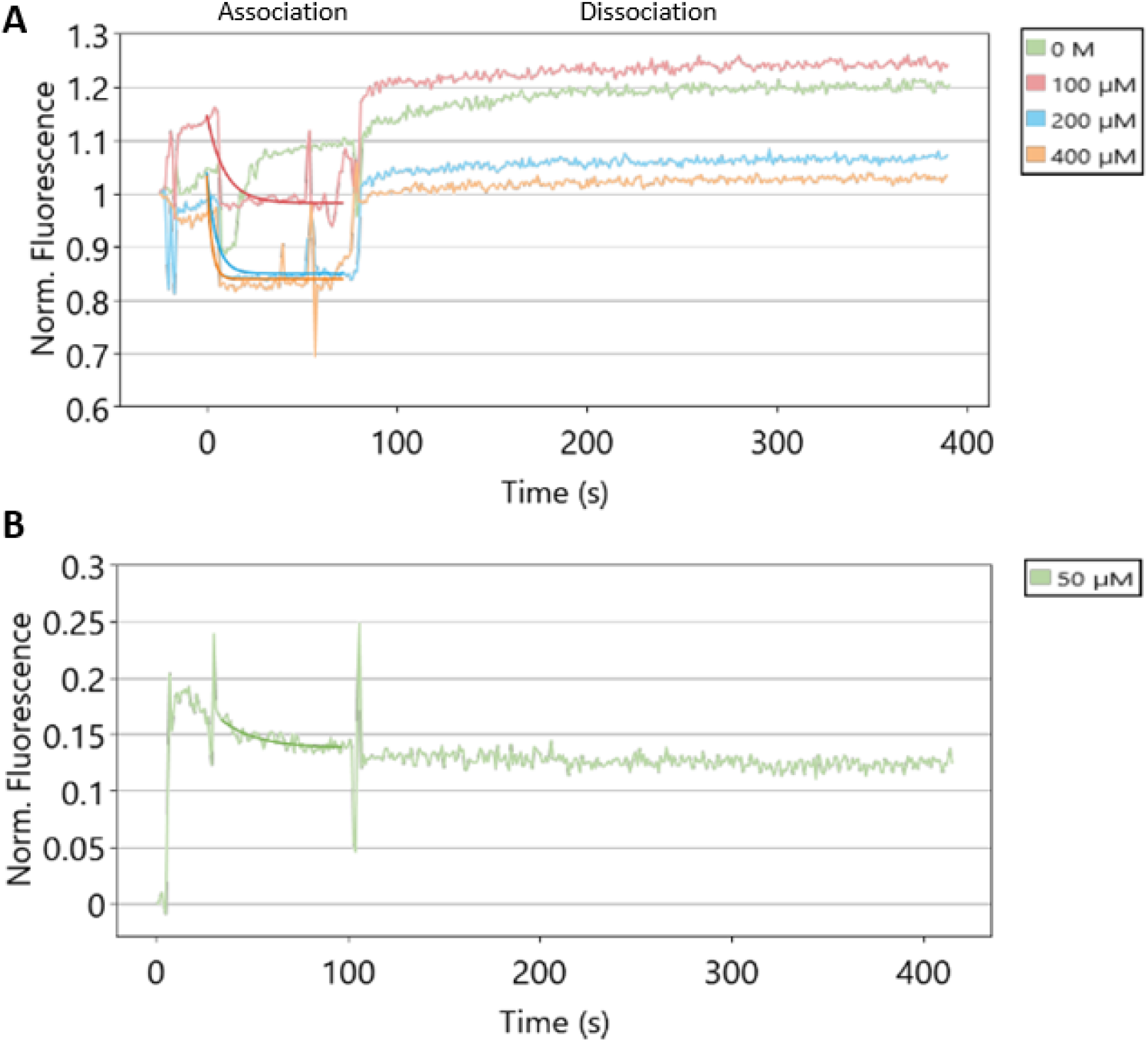
Data analysis using switchANALYSIS. Kinetic characterization of AA8AC40 binding to Sirt3 using dynamic mode. Concentration depending decrease in dynamic response upon association and dissociation. Global mono-exponential fit to determine rate constants and affinity. **(A)** in the presence of cofactors, OAADPr + QPKK-NH_2_; **(B)** in the absence of cofactors.

**Table 3.** Overview of the obtained rate constants and affinity of compound AA8AC40 under different conditions.

|  | AA8AC40 |  |
| --- | --- | --- |
|  | Sirt3+OAADPr + QPKK-NH<sub>2</sub> | Sirt3 |
| $K_{on}$, M<sup>-1</sup>s<sup>-1</sup> | 9.60 ± 1.50 E+2 | 1.02 ± 0.32 E+3 |
| $K_{off}$, s<sup>-1</sup> | 1.21 ± 0.06 E-2 | 4.63 ± 3.24 E-3 |
| $K_d$, μM | 12.6 ± 2.1 | 4.56 ± 3.50 |

## 3. Conclusion

In this study, we have produced and selected a 3880 member DNA encoded library against Sirt3 in the presence of various cofactors (Carba-NAD or OAADPr). The several identified hits were tested off-DNA by activity assay, and we have reported various novel inhibitors under steady state conditions and several activators under non-steady state conditions.

Our best activator AA8AC40 has a similar potency to Honokiol (∼107% at 10µM) and a binding affinity in the low micromolar range (4.6 µM). The docking studies suggest that our modulators bind to an atypical site of Sirt3. Further studies should be useful to clearly identify the binding site and the modulation mechanism of these compounds.

## 4. Materials and Methods

### 4.1. General

For Oligoconjugate derivatives: HPLC were performed on a Waters Oligonucleotides BEH C18 50×4.6 mm, 2.5 µm column using 100 mM TEAA, pH 7 and MeCN in gradient mode. A MALDI-TOF/TOF UltrafleXtreme mass spectrometer (*Bruker Daltonics, Bremen*) was used to check the masses of oligoconjugated amino acids. Mass spectra were obtained in linear positive ion mode. The laser intensity was set just above the ion generation threshold to obtain peaks with the highest possible signal-to-noise (S/N) ratio without significant peak broadening.

For Off-DNA compounds: UPLC/MS were performed on a Waters Acquity H-Class with QDa and PDA detectors with an Acquity UPLC BEH C18 50×2.1 mm – 1.7 µm using water/0.01% formic acid and Acetonitrile/0.01% formic acid in gradient mode. NMR were performed on a Spinsolve 80 Carbon Benchtop Spectrometer from *Magritek Gmbh*.

### 4.2. DEL Library synthesis

#### Coupling Reactions of 20 Fmoc-Protected Amino Acids

To a reaction volume of 310 µL containing 70% (vol/vol) DMSO/water, compounds were added to the respective final concentrations: Fmoc-protected amino acids DMSO solution, 4 mM; N-hydroxysulfosuccinimide in DMSO, 10 mM; N-ethyl-N9-(3-dimethylaminopropyl) carbodiimide in DMSO, 4 mM; aqueous triethylamine hydrochloride solution (pH 9.0), 80 mM; oligonucleotide aqueous solution, 50 mM, (59-GGA GCT TGT GAA TTC TGG XXX XXX GGA CGT GTG TGA ATT GTC-39, XXX XXX unambiguously identifies the individual Fmoc-protected amino acid compound). All coupling reactions were stirred overnight at 25 °C; residual activated species were then quenched and simultaneously Fmoc-deprotected by addition of piperidine (500 mM in DMSO). The reactions were then purified by HPLC, and the desired fractions were dried under reduced pressure and redissolved in 100 µL of water. Four nanomoles of each DNA– compound conjugate was pooled to generate a 20-member DNA-encoded sub-library.

#### Coupling Reactions of 200 Carboxylic Acids

To a reaction volume of 310 µL, containing 70% (vol/vol) DMSO/water, compounds were added to the respective final concentrations: DMSO-dissolved carboxylic acid, 4 mM; N-hydroxysulfosuccinimide in DMSO, 10 mM; N-ethyl-N9-(3-dimethylaminopropyl)-carbodiimide in DMSO, 4 mM; aqueous triethylamine hydrochloride (pH9.0), 80 mM; DNA-oligonucleotide sublibrary pool, 1.5 mM. All coupling reactions were stirred overnight at 25 °C; residual activated species were then quenched by addition of 50 mL of Tris-Cl buffer, 500 mM (pH 9.0). The mixture was allowed to quantitatively precipitate by sequential addition of 25 mL of 1 M acetic acid, 12.5 mL of 3 M sodium acetate buffer (pH 4.7), and 500 mL of ethanol followed by 2-h incubation at -20 °C. The DNA was centrifuged, and the resulting oligonucleotide pellet was washed with ice-cold 90% (vol/vol) ethanol and then dissolved in 60 µL of water. Test coupling reactions were also performed with the reaction conditions described above; using model 42-mer 5’-Fmoc-deprotected amino acid oligonucleotide conjugates and model carboxylic acids. Concentrations were measured by UV spectrophotometry.

#### Polymerase Klenow Encoding of 194 Carboxylic Acid Reactions

To a reaction volume of 50 µL, reagents were added to the respective final concentrations: pool 20 conjugates 320 nM, 44-mer oligo-nucleotide (59-GTA GTC GGA TCC GAC CAC XXXX XXXX GAC AAT TCA CAC ACG TCC-39, XXXX XXXX unambiguously identifies the individual carboxylic acid compound, IBA) 600 nM, Klenow buffer (cat. no. B7002S; NEB), dNTPs (cat. no.11969064001; Roche), 0.5mM, Klenow Polymerase enzyme (cat. no. M0210L; NEB), 5 units. The Klenow polymerization reactions were incubated at 37 °C for 1 h and then purified on ion-exchange cartridges (cat. no. 28306; Qiagen). The 194 purified reactions were dissolved in 50 µL of water each and pooled to generate the 3,880 members library (DEL3880) to a final total oligonucleotide concentration of 100 nM.

### 4.3. DNA encoded library selection against SIRT-3

A N-terminal His tagged Sirt3(118-399) from Abcam was used for all the DEL experiments. Before selection, the activity of Sirt3 was checked by a fluorometric assay kit (Abcam). This construct contains NAD binding pocket and the C-terminal.

#### Selection of binding molecules to Sirt3

Affinity-mediated selection of the binding molecules against recombinant Sirt3 was done by incubating 20 nM of the DEL library with 50 µg of Sirt3 in 400 µl HDACT buffer (Enzo life sciences) containing either Carba-NAD,100 µM (Dalton pharma, Toronto) or OAADPr 100 µM (Santa-Cruz biotechnology), peptide substrate QPKK(Ac),50 µM (Proteogenix), Herring sperm DNA (0.32 mg/ml). This mix was added to 50 µl pre-washed His-Pur magnetic Nickel beads (Thermofisher Scientific) and incubated for 1 h at 4 °C on a rotor. Two different samples were prepared, one containing Sirt3 (target selection) and the other without Sirt3 (no target selection) that served as a negative control. Following selection, the supernatant was discarded, and the beads were washed (4*400 µl HDACT buffer) to remove any unbound molecules. Elution was carried out in 110 µL of HDACT buffer by incubating the beads at 75 °C for 15 min for both the samples. The encoding oligonucleotides in the elution fraction were PCR amplified by Phusion polymerase for 15 cycles at 30 s at 98 °C, denaturation 15 s at 98 °C, annealing 60 °C in 15 s and extension 1 min at 72 °C. Following which the PCR products were purified by PCR cleanup kit (NEB) and a second PCR was done with primers containing the READ1, READ2 and adapter sequences required to support clustering and subsequent single-read 150bp sequencing on an Illumina iseq100. A total of 8 million sequence reads were analyzed The Q30 values of both Read1 and Read2 were above 90%.

All DNA sequences with a phred score above 30 were taken for analysis. Firstly, the data was sorted according to the “sample barcode” that was added in the PCR post screening. This sample barcode allowed us to pool all the different screening samples and sequence all at once. After sorting, the unique barcode sequences corresponding to the 20 amino acids and 194 carboxylic acids were located and counted in each screening sample. In order to remove amplification bias, the raw reads were normalized with respect to the total number of reads in each sample generating a frequency value for each compound in the target selection sample and the no target selection sample. The enrichment for each compound was calculated as the ratio of their frequency in target selection sample to the no target selection sample. The frequency of the highly enriched compounds was verified in the native library to make sure than enrichment was indeed due to binding to Sirt3 and not merely because of a high starting concentration. 3D graphs were generated by an in-house program written in python.

### 4.4. Binders off-DNA syntheses

#### Off-DNA synthesis of the binding molecules

##### Synthesis of 2-[(2-amino-6-tricyclo[9.4.0.03,8]pentadeca-1(11),3(8),4,6,12,14-hexaenyl)oxy]-N-butylacetamide (Scaffold 1)

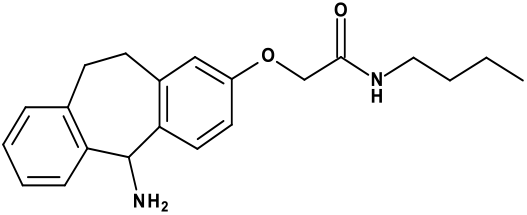

To a solution of Ramage Linker or Fmoc-Suberol (5 g, 9.9 mmol) in DMF (50 ml), was added HBTU (5.6 g, 14.85 mmol) and NaHCO_3_ (1.66 g, 19.8 mmol). After stirring at RT for 0.5 h, n-butylamine was added (0.86 g, 11.86 mmol). The reaction mixture was stirred at RT for 2 h then quenched by adding 380 ml of water. A precipitate was formed, stirred at RT for 1hr then filtered and washed with water (250 ml x 2). The beige solid was dried in vacuum oven overnight and then directly engaged into the next step. To a solution of the obtained beige powder in methanol (100 mL) at 0-5 °C, was added piperidine (13.20 ml, 133.7 mmol). The mixture was slowly allowed to warm at RT and stirred at RT for 1 hr. THF (200 ml) was added to improve the solubility and the mixture was stirred at RT overnight. The mixture was concentrated under reduced pressure. The crude product was purified by flash column chromatography (DCM/MeOH gradient) to afford **scaffold 1** (3 g, 85% yield) as a beige solid. LCMS Calculated for C_21_H_26_N_2_O_2_, 338.2, Observed [M+Na]^+^ 361.

1H NMR (80 MHz, DMSO-*d6*) δ 8.33 – 7.75 (m, 1H), 7.55 – 7.24 (m, 2H), 7.11 (d, J = 2.7 Hz, 3H), 6.70 (m, 2H), 5.40 (s, 1H), 4.38 (s, 2H), 3.12 (m, 7H), 1.76 – 1.00 (m, 5H), 0.88 (m, 3H)

##### Synthesis of (2R)-2-amino-N-butyl-3-(4-chlorophenyl)propanamide) (Scaffold 2)

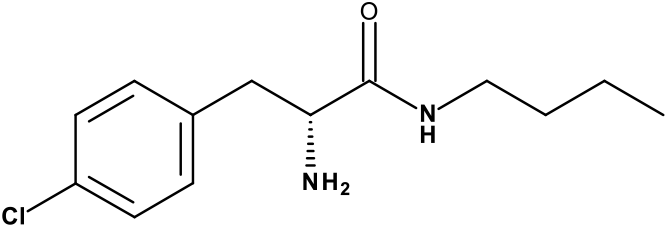

To a solution of (2S)-1-(4-chlorophenyl)-*N*-methyl-propan-2-amine (4.9 g, 11.7 mmol) in THF (45 ml), was added DMT.MM (3.88 g, 14.03 mmol) and DIEA (4.07 ml, 23.38 mmol). After stirring at RT for 15 mins, *n*-butylamine was added (0.94 g, 12.85 mmol). The reaction mixture was stirred at RT overnight. The mixture was concentrated under reduced pressure then 150 ml of water was added. A precipitate was formed, it was filtered and washed with water (250 ml x 2). The white solid was dried in vacuum oven at 40 °C overnight and then directly engaged into the next step.

To a solution of the obtained white powder in methanol (100 mL) at 0 °C, was added pyridine (15.6 ml, 157.8 mmol). The mixture was slowly allowed to warm at RT and stirred at RT overnight.

The mixture was concentrated under reduced pressure. The crude product was roughly purified by flash column chromatography (DCM/MeOH gradient) to afford scaffold 1 (1.35 g, 39% yield) as a yellow solid. The compound purity was not good enough to provide a reliable NMR description, but the product was used in the downstream chemistry without further purification.

LCMS Calculated for C_13_H_19_ClN_2_O, 254.1, Observed [M+H]^+^ 255.

##### General procedure for peptidic couplings on Scaffold 1

With carboxylic acids:

AC71: S-(Thiobenzoyl)thioglycolic acid (**1**)

AC92: Ethacrynic acid (**2**)

AC40: 4-[4-(1-Hydroxyethyl)-2-methoxy-5-nitrophenoxy]butyric acid (**3**)

To a solution of each carboxylic acid (**1**: 113 g, **2**: 161.3 g, **3**: 159.3 g) in 2mL of DMF at RT, were added HBTU (1,5 eq.) then DIEA (2 eq.) successively. The reactions were stirred at RT for 30 min then **Scaffold 1** (150 mg) was added to each reaction. The reactions were stirred at RT overnight. In each reaction, 10 ml of water were added. Precipitates were formed and filtered then washed with water.

Products were purified by flash column chromatography (DCM/MeOH gradient) to give:

**AA8AC71**: (120 mg, 51% yield) as a pink solid.

LCMS Calculated for **AA8AC71** C_30_H_32_N_2_O_3_S_2_, 532.1, Observed [M+Na]^+^ 555/557. 1H NMR (80 MHz, DMSO-*d6*) δ 9.33 (d, J = 7.8 Hz, 1H), 7.95 (d, J = 6.2 Hz, 3H), 7.68 – 7.23 (m, 5H), 7.19 (s, 3H), 6.75 (d, J = 6.9 Hz, 2H), 6.21 (d, J = 7.6 Hz, 2H), 4.38 (d, J = 6.2 Hz, 4H), 3.33 – 2.68 (m, 5H), 1.73 – 1.08 (m, 4H), 1.10 – 0.42 (m, 3H)

13C NMR (20 MHz, DMSO*-d6*) δ 167.37, 164.30, 157.01, 143.98, 140.03, 139.24, 138.64, 132.98, 131.80, 129.91, 129.76, 129.24, 128.75, 127.76, 127.37, 126.40, 125.87, 116.22, 111.86, 66.99, 55.51, 41.30, 39.09, 37.88, 32.03, 31.56, 31.17, 19.46, 13.61

**AA8AC92**: (135 mg, 49% yield) as a white solid.

LCMS Calculated for **AA8AC92** C_34_H_36_Cl_2_N_2_O_5_, 622, Observed [M+H]^+^ 623, [M+Na]^+^ 645/647. 1H NMR (80 MHz, DMSO*-d6*) δ 9.03 (d, J = 7.7 Hz, 1H), 7.95 (t, J = 5.6 Hz, 1H), 7.53 – 7.04 (m, 7H), 6.97 – 6.57 (m, 2H), 6.24 (d, J = 7.7 Hz, 1H), 6.06 (s, 1H), 5.54 (s, 1H), 4.86 (s, 2H), 4.41 (s, 2H), 3.29 – 2.87 (m, 5H), 2.45 – 1.90 (m, 1H), 1.80 – 0.61 (m, 12H)

13C NMR (20 MHz, DMSO*-d6*) δ 195.09, 167.35, 165.46, 157.04, 155.59, 149.36, 139.99, 139.09, 138.60, 132.35, 131.62, 129.97, 129.27, 127.73, 127.37, 125.91, 121.06, 116.25, 111.87, 67.73, 66.99, 54.87, 37.89, 32.04, 31.59, 31.18, 22.94, 19.46, 13.61, 12.33

**AA8AC40**: (50 mg, 18% yield) as a light yellow solid.

LCMS Calculated for **AA8AC40** C_34_H_41_N_3_O_8_, 619.2, Observed [M+Na]^+^ 642.

1H NMR (80 MHz,DMSO*-d6*) δ 8.89 (d, J = 7.9 Hz, 1H), 7.97 (s, 1H), 7.79 – 7.02 (m, 7H), 6.70 (d, J = 7.7 Hz, 2H), 6.29 (d, J = 7.9 Hz, 1H), 5.48 (d, J = 4.5 Hz, 1H), 5.26 (t, J = 5.6 Hz, 1H), 4.39 (s, 2H), 4.17 – 3.77 (m, 5H), 3.24 – 2.85 (m, 6H), 2.34 (sl, 1H), 2.21 – 1.76 (m, 2H), 1.57 – 1.14 (m, 8H), 0.87 (d, J = 6.2 Hz, 3H)

13C NMR (20 MHz, DMSO*-d6*) δ 175.58, 170.13, 167.04, 156.47, 153.07, 145.91, 139.50, 138.50, 138.11, 137.66, 132.04, 129.48, 128.27, 126.78, 125.43, 115.80, 111.46, 108.73, 107.96, 67.86, 66.65, 63.53, 55.68, 37.53, 31.65, 31.22, 31.06, 30.82, 24.79, 24.43, 19.10, 13.25

##### General procedure for peptidic couplings on Scaffold 2

With carboxylic acids:

AC71: S-(Thiobenzoyl)thioglycolic acid (**1**)

AC92: Ethacrynic acid (**2**)

AC78: 6-Acetylthiohexanoic acid (**4**)

AC02Bis: 4-[(2,4-Dioxo-1,3-thiazolidin-5-ylidene)methyl]benzoic acid (**5**)

To a solution of each carboxylic acid (**1:** 211.9 g, **2:** 180.4 g, **3:** 161.7 g, **4:** 257.7 g) in 3 mL of THF at RT, were added DMT.MM (1.2 eq.) then DIEA (2 eq.) successively. The reactions were stirred at RT for 30 min then 2 mL of a solution of **Scaffold 2** (908 mg in 10 mL of THF) was added to each reaction. The reactions were stirred at RT overnight.

In each reaction, a precipitate was formed. The insoluble was filtered and rinsed with THF (10 ml x 2). the filtrate was concentrated under vacuum to give colored powders. Products were purified by flash column chromatography (DCM/MeOH gradient) to give:

**AA12AC71**: (90 mg, 28 % yield) as a pink solid.

LCMS Calculated for **AA12AC71** C_22_H_25_ClN_2_O_2_S_2_, 448.1, Observed [M+H]^+^ 449/451.

1H NMR (80 MHz,DMSO*-d6*) δ 8.59 (d, J = 8.3 Hz, 1H), 7.93 (dd, J = 6.1, 2.0 Hz, 3H), 7.74 – 7.36 (m, 3H), 7.28 (s, 4H), 4.53 (q, 1H),4.18 (s, 2H), 3.19 – 2.74 (m, 4H), 1.18 (dd, J = 13.2, 4.3 Hz, 4H), 0.88 (d, J = 6.2 Hz, 3H)

13C NMR (20 MHz, DMSO-d6) δ 166.61, 161.75, 140.52, 133.24, 129.61, 127.64, 125.36, 124.59, 123.02, 50.89, 37.72, 34.79, 27.66, 16.05, 10.24

**AA12AC92**: (70 mg, 18 % yield) as a white solid.

LCMS Calculated for **AA12AC92** C_26_H_29_Cl_3_N_2_O_4_, 538,1, Observed [M+H]^+^ 539/541.

1H NMR (80 MHz, DMSO*-d6*) δ 8.48 – 7.87 (m, 3H), 7.49 – 7.10 (m, 4H), 6.81 (d, J = 8.7 Hz, 1H), 6.07 (s, 1H), 5.55 (s, 1H),5.08 – 4.20 (m, 3H), 3.00 (t, J = 6.9 Hz, 4H), 2.29 (d, J = 7.4 Hz, 2H), 1.51 – 0.21 (m, 10H).

13C NMR (20 MHz, DMSO*-d6*) δ 195.01, 169.95, 166.23, 155.34, 149.35, 136.52, 132.38, 131.08, 129.43, 128.02, 127.29, 121.08, 111.59, 67.46, 53.35, 38.23, 31.06, 22.97, 19.46, 13.63, 12.37

**AA12AC78**: (170 mg, 56 % yield) as a white solid.

LCMS Calculated for **AA12AC78** C_21_H_31_ClN_2_O_3_S, 426.1, Observed [M+H]^+^ 427/429. 1H NMR (80 MHz,CDCl_3_) δ 7.44 (dd, J = 8.4, 3.4 Hz, 4H), 6.69 (d, J = 8.1 Hz, 1H), 6.40 (t, 1H), 4.83 (q, J = 7.5 Hz, 1H), 3.15 (tt, J = 13.1, 6.2 Hz, 6H), 2.51 (s, 3H), 2.43 – 2.02 (m, 2H), 1.99 – 1.23 (m, 10H), 1.20 – 0.92 (m, 3H)

13C NMR (20 MHz, DMSO*-d6*) δ 196.02, 172.92, 170.78, 135.44, 132.93, 130.80, 128.77, 54.50, 39.35, 38.26, 36.29, 31.50, 30.76, 29.37, 28.92, 28.32, 25.12, 20.06, 13.79

**AA12AC02Bis**: (96 mg, 29 % yield) as a light yellow solid.

LCMS Calculated for **AA12AC02Bis** C_24_H_24_ClN_3_O_4_S, 485.1, Observed [M+H]^+^ 486/488. 1H NMR (80 MHz, DMSO-*d6*) δ 12.64 (s, 1H), 8.70 (d, J = 8.3 Hz, 1H), 7.79 (dt, J = 17.0, 8.2 Hz, 6H), 7.33 (s, 4H), 4.66 (q, J = 8.4 Hz, 1H), 3.18 – 2.83 (m, 4H), 1.60 – 1.05 (m, 4H), 1.00 – 0.64 (m, 3H)

13C NMR (20 MHz, DMSO-*d6*) δ 168.80, 165.95, 165.51, 163.53, 135.49, 133.82, 133.15, 129.16, 128.72, 127.86, 126.40, 126.17, 123.56, 53.01, 34.93, 29.28, 17.63, 11.81

### 4.5. Activity assays of hit compounds

#### Chemicals and Reagents

MnSOD (KGELLEAI-(KAc)-RDFGSFDKF) was synthesized by GenScript (Piscataway, NJ). FdL2 (QPKK^AC^-AMC) peptide, also called p53-AMC peptide, was purchased from Enzo Life Sciences (Farmingdale, NY). Carba-NAD was synthesized by Dalton Pharma (Toronto, ON). The hSirt3^118-399^ was purchased from Xtal Biostructures (Natick, MA).

#### Effect of hit compounds on hSirt3^1118-399^ deacetylation activity

##### HPLC assay using native peptide

Enzymatic reactions included either 1 mM NAD^+^ and 50 µM MnSOD peptide or 50 µM NAD^+^ and 600 µM peptide substrate in presence of hit compounds (10 µM), in a buffer (50mM TRIS-HCl, 137mM NaCl, 2.7mM KCl, and 1mM MgCl_2_, pH 8.0 and 5% DMSO). The reaction was initiated by adding 5U hSirt3^118-399^ and incubated at 37 °C for 30 minutes. The reaction was terminated by 2%TFA. The peptide product and substrate were resolved using HPLC. Solvent A was composed of 90% HPLC grade water and 10% acetonitrile with 0.02% (v/v) TFA. Solvent B was composed of acetonitrile with 0.05% (v/v) TFA. A linear gradient was performed for 20 min from 0% B to 51% (v/v) B. Solvent B percentage was increased to 100% within 5 minutes. Then returned to the starting conditions (0% B) within 5 min. Solvent A percentage (100%) was maintained at 100% for 5 additional minutes (36 min total run time).

##### Fluorescence-based assay using Fluorolabeled Peptide

The modulation effect of hits compounds for Sirt3^118-399^ deacetylation activity was determined using FdL2 peptide. The enzymatic reactions were carried out as described above. The reaction was terminated by adding the 1x developer 2mM NAM solution and measured the fluorescence on TECAN microplate reader. The raw data were fitted to the defined model equations using GraphPad Prism (GraphPad Software, Inc, CA).

### 4.6. Binding affinity measurement by switchSENSE®

Binding assays were performed with switchSENSE® technology using a DRX^2^ device (Dynamic Biosensors GmbH) at the Institute of integrative Biology of the Cell (I2BC – University Paris Saclay, CEA, CNRS). Standard multipurpose biochips MPC-48-1-R1-S or MPC2-48-2-G1R1-S were used for measurements (Dynamic Biosensors GmbH). They were used according to the manufacturer’s instructions. PE40 (10 mM Na_2_HPO_4_/NaH_2_PO_4_, 10 mM Na_2_HPO_4_/NaH_2_PO_4_, 40 mM NaCl, 0.05 % Tween 20, 50 μM EDTA, 50 μM EGTA) was used as running and auxiliary buffer as recommended by the manufacturer. Complementary DNA strands (such as cNL-B48) were purchased from Dynamic Biosensors GmbH, as well as regeneration solution and glass vials. All consumables were used according to the manufacturer’s instructions. Measurement protocols were established using the switchBUILD software and carried out using switchCONTROL software (both provided by Dynamic Biosensors GmbH). Each flow channel was passivated prior to measurements using a thiol-containing passivation solution (Dynamic Biosensors GmbH). Afterward, the electrode that showed the highest fluorescence amplitude in the chip status test was selected for further measurements. One flow channel was typically used for up to four measurement cycles. Immobilization of the DNA constructs was done by conjugate hybridization using a concentration of 500 nM for an on-chip hybridization time of 320 s. Interaction measurements were carried out in static mode. Binders were dissolved in DMSO and diluted with HDAC buffer to 3 concentrations ranged from 100 to 400 µM in 5%DMSO/HDAC solution. Co-factors (Carba-NAD/QPKKAc or OAADPR/QPKK-NH_2_) were added at 100 µM/50 µM concentration when specified. All data were acquired with the system temperature set to 25 °C. The association process was done at a flow speed of 200 μL min^−1^ for 100 s. The dissociation volume was done at a flow speed of 1000 μL min^−1^ for 285 s. After measurements, a standby routine was performed to store the biochip with double stranded DNA.

### 4.7. Docking studies

#### MD simulation

Computational studies were performed using MOE (Molecular Operating Environment from Chemical Computing Group Inc., Montreal, PQ) version MOE 2020.09 on computers with quad-core Intel x86-64 (Intel Core i7- 8th Generation) processors, 16 GB of RAM and running Windows 10 Pro. MMFF94x force field and solvation, as implemented in MOE, was used for all computational studies.^22–24^

#### Ligand docking procedure

We used two methods to define the ligand-binding site in receptor protein structures. The first one, used for 4BVG, defined the site using the co-crystallized ligand (EX-527). The second, employed for 4FVT and 4BVG, utilized the ‘Site Finder module’ in MOE to create and place dummy atoms into potential ligand-binding sites for both structures. This step was performed after removing all molecules of minor compounds present within receptor structures such as sulphate ions, glycerol, 1,2-ethanediol etc. Fortuitously, the major site identified by second method for 4BVG was almost identical to the one defined by co-crystallized ligand. Hence it was assumed that the main putative (and internal) ligand-binding site identified in 4FVT was the best candidate for docking studies in that receptor structure.

Energy-optimized models of selected compounds were sequentially docked into these sites by ‘Dock’ application in MOE; default settings for parameters using the ‘Rigid Receptor’ protocol was employed for all docking studies. Top poses which exhibited reproducibility (RMSD < 0.2 Å) in three independent docking simulations at a given site were the most optimal poses for that site. The main settings for ligand docking to were as follows: Docking Protocol = Rigid Receptor; Placement = Triangle Matcher; Rescoring = London dG; Retain = 100, Refinement = Forcefield; Rescoring = GBVI/WSA dG; Retain = 100 top poses. The default settings for docking parameters using ‘Rigid Receptor’ protocol was found to be satisfactory for docking of Honokiol and EX-527, which was used as test ligands. The top scoring pose for ligands from each docking run was used for calculating the approximate binding energy and other inputs for computation of the final docking scores.

#### Calculation of ligand binding energy

The binding energy for top docked poses of each ligand at each site was calculated using an updated version of a published method.^25,26^ It involved: calculating energies of the docked receptor-ligand complex (Ecpx); the unbound ligand in solution (Elig) and unbound solvated rigid receptor from the complex (Eprot) for use in the equation: Ebind = Ecpx - (Elig + Eprot). The top selected docked pose of each ligand, obtained from three independent docking runs, was used for calculation of the approximate ligand-binding energy. Optimization of semi-rigid (fixed heteroatoms) receptor-ligand complex was conducted under distance-dependent dielectric conditions, the final step being performed under solvation, using a regime under which only hydrogens in the ligand and cavity could undergo further structural optimization.

Binding energies calculated using molecular mechanics should always be viewed as approximations of the actual binding energy since any semi-rigid receptor approximation almost always underestimates effects of induced fit, whereas a fully flexible receptor approximation usually overestimates it. Hence, actual binding energy would be expected to lie between the rigid and flexible receptor values. The methodology used for the calculation of receptor buriedness for top docked (and energy optimized) ligand poses, as well as enumeration of their hydrogen bond interactions with residues in the binding site have been described previously; and the code is available upon request.^26^

## Acknowledgement

We would like to thank Dr. Paloma Fernandez-Varela and Dr. Jean-Baptiste Charbonnier from Université Paris-Saclay, CEA, CNRS, Institute for Integrative Biology of the Cell (I2BC), (Gif-sur-Yvette, France) for their contribution with the binding affinity assays.

This research did not receive any specific grant from funding agencies in the public, commercial, or not-for-profit sectors.

## Declaration of interests

The authors declare that they have no known competing financial interests or personal relationships that could have appeared to influence the work reported in this paper.

## Notes

### Competing Interest Statement

The authors have declared no competing interest.

